# Genomics and Phenomics Enabled Prebreeding Improved Early-Season Chilling Tolerance in Sorghum

**DOI:** 10.1101/2022.10.31.514536

**Authors:** Sandeep Marla, Terry Felderhoff, Chad Hayes, Ramasamy Perumal, Xu Wang, Jesse Poland, Geoffrey P. Morris

## Abstract

In temperate climates, earlier planting of tropical-origin crops can provide longer growing seasons, reduce water loss, suppress weeds, and escape post-flowering drought stress. However, chilling sensitivity of sorghum, a tropical-origin cereal crop, limits early planting and over 50 years of conventional breeding has been stymied by coinheritance of chilling tolerance (CT) loci with undesirable tannin and dwarfing alleles. In this study, phenomics and genomics-enabled approaches were used for prebreeding of sorghum early-season CT. Uncrewed aircraft systems (UAS) high-throughput phenotyping platform tested for improving scalability showed moderate correlation between manual and UAS phenotyping. UAS normalized difference vegetation index (NDVI) values from the chilling nested association mapping population detected CT QTL that colocialized with manual phenotyping CT QTL. Two of the four first-generation KASP molecular markers, generated using the peak QTL SNPs, failed to function in an independent breeding program as the CT allele was common in diverse breeding lines. Population genomic *F*_ST_ analysis identified SNP CT alleles that were globally rare but common to the CT donors. Second-generation markers, generated using population genomics, were successful in tracking the donor CT allele in diverse breeding lines from two independent sorghum breeding programs. Marker-assisted breeding, effective in introgressing CT allele from Chinese sorghums into chilling-sensitive US elite sorghums, improved early-planted seedling performance ratings in lines with CT alleles by up to 13–24% compared to the negative control under natural chilling stress. These findings directly demonstrate the effectiveness of high-throughput phenotyping and population genomics in molecular breeding of complex adaptive traits.

## INTRODUCTION

Climate stress is a major limiting factor for crop production. To minimize the yield losses by adverse weather patterns for providing global food security, breeding food crops with abiotic stress tolerance is critical (Mickelbart et al., 2015). Conventional breeding efforts to increase climate resilience in food crops were slow due to the complex genetic architecture of abiotic stress tolerance (Collins et al., 2008; Bernardo, 2016). Further complicating conventional breeding for abiotic stress tolerance are spatiotemporal variability of environmental conditions in targeted environments between years, lack of reliable phenotyping for abiotic stress tolerance traits, and poor scalability of field trials due to limited phenotyping resources (Hickey et al., 2019; Varshney et al., 2021). Given that conventional breeding was not effective in developing abiotic stress tolerance, there is a need to utilize modern breeding tools in combination with modern phenotyping platforms in abiotic stress tolerance breeding.

Substantial advances in next-generation sequencing technologies over the past two decades established genomics-assisted breeding as an alternate solution to conventional breeding for complex traits (Sasaki, 2005; Schnable et al., 2009; Abberton et al., 2016; McCormick et al., 2018; Tao et al., 2019; Varshney et al., 2021). Despite the advances in genomics-assisted breeding, breeding for complex traits mostly failed in public breeding due to weak association between the phenotype and the genomic region (i.e. trait-to-QTL correlation), the QTL and the marker used for selecting the trait (i.e. the QTL-to-marker correlation), and failure of QTL to function in diverse genetic backgrounds (Cobb et al., 2018). Increasing mapping power and resolution with multiparental populations and high density markers can improve trait-to-QTL associations for adaptive traits (Yu et al., 2008; Bouchet et al., 2017; Gage et al., 2020). To improve QTL-to-marker associations across diverse genetic backgrounds in a breeding program, population genomics approaches can be used to identify fixed alleles in locally adapted germplasm, that are in linkage disequilibrium (LD) with the trait of interest (Muleta et al., 2019). Therefore, there is a need for public breeding programs to integrate high-resolution mapping and population genomics approaches to develop abiotic stress tolerance in food crops.

Sorghum (*Sorghum bicolor*) is a tropical-origin C4 cereal crop that is sensitive to chilling temperatures (0–15°) (Taylor & Rowley, 1971; Lyons, 1973). Developing chilling-tolerant sorghum hybrids could enable early-planting in temperate regions, increase growing period by 20%, reduce evaporation during fallow periods, reduce risk of terminal droughts, and increase 7– 11% grain yield (Raymundo et al., 2021). Chinese sorghums, introduced from Africa into China ~800 years ago and adapted to chilling temperatures, were a good source for chilling tolerance breeding (Stickler et al., 1962; Salas Fernandez et al., 2014; Marla et al., 2019). Coarse resolution mapping, fewer recombinant inbred lines (RILs) and low marker density, with US chilling-sensitive and Chinese chilling-tolerant biparental families identified CT loci linked with undesirable tannin and plant height alleles (Knoll et al., 2007; Burow et al., 2010; Ostmeyer et al., 2020). High-resolution CT mapping with a chilling nested association mapping (NAM) population detected precise co-localization of CT alleles with undesirable grain tannin alleles at *Tan1* (*Tannin1*) and *Tan2* (*Tannin2*) genes, and tall plant alleles at *Dw1* (*Dwarf1*) and *Dw3* (*Dwarf3*) genes (Marla et al., 2019). Coinheritance of CT loci with undesirable tannin and dwarfing alleles combined likely stymied conventional chilling tolerance breeding for over 50 years.

New genome-to-phenome (G2P) approaches offer potential solutions for complex trait breeding that was hindered by low-resolution mapping, poor scalability, and complex phenotyping for abiotic stress tolerance (Cooper et al., 2014; Kole et al., 2015; Abberton et al., 2016; Varshney et al., 2021). We hypothesized G2P-enabled strong trait-to-QTL and QTL-to-marker association improves sorghum early-season chilling tolerance. Based on this hypothesis, we predicted elite US sorghum lines with CT alleles, generated using marker-assisted breeding, will exhibit higher early-season seedling emergence count and seedling vigor ratings compared to the negative controls. Joint linkage mapping (JLM) of UAS phenomic data on the chilling NAM population and fixation index (*F*_ST_) population genomic analysis generated Kompetitive Allele Specific PCR (KASP) genotyping markers for precisely introgressing CT alleles from Chinese lines into chilling-sensitive US elite lines. Segregating families and sorghum hybrids carrying CT alleles, generated using marker-assisted breeding, contained higher seedling performance ratings under natural chilling stress. Overall, the study demonstrates the power of combining phenomics and genomics-enabled prebreeding for complex adaptive traits.

## RESULTS

### UAS phenomics validated CT QTL detected with manual seedling vigor ratings

To evaluate the potential of UAS HTP platforms for chilling tolerance mapping, UAS phenotyping was conducted on two sorghum early-planted field trials, AB17 and MN17 (Figure 1a). Manual phenotyping for early-season seedling performance ratings in AB17 field trial required ~5 h for each trait. As manual phenotyping was time-consuming, only three seedling vigor ratings were collected manually for each field trial. In contrast, UAS phenotyping enabled capturing NDVI values several times during the seedling stage. Correlation analysis between AB17 UAS NDVI values and their corresponding three manual SV ratings demonstrated an intermediate correlation (0.58, 0.53, and 0.54) (Figure 1b). Similarly, MN17 NDVI value and its corresponding manual SV rating showed a moderately high correlation (0.67).

**Figure 1:**
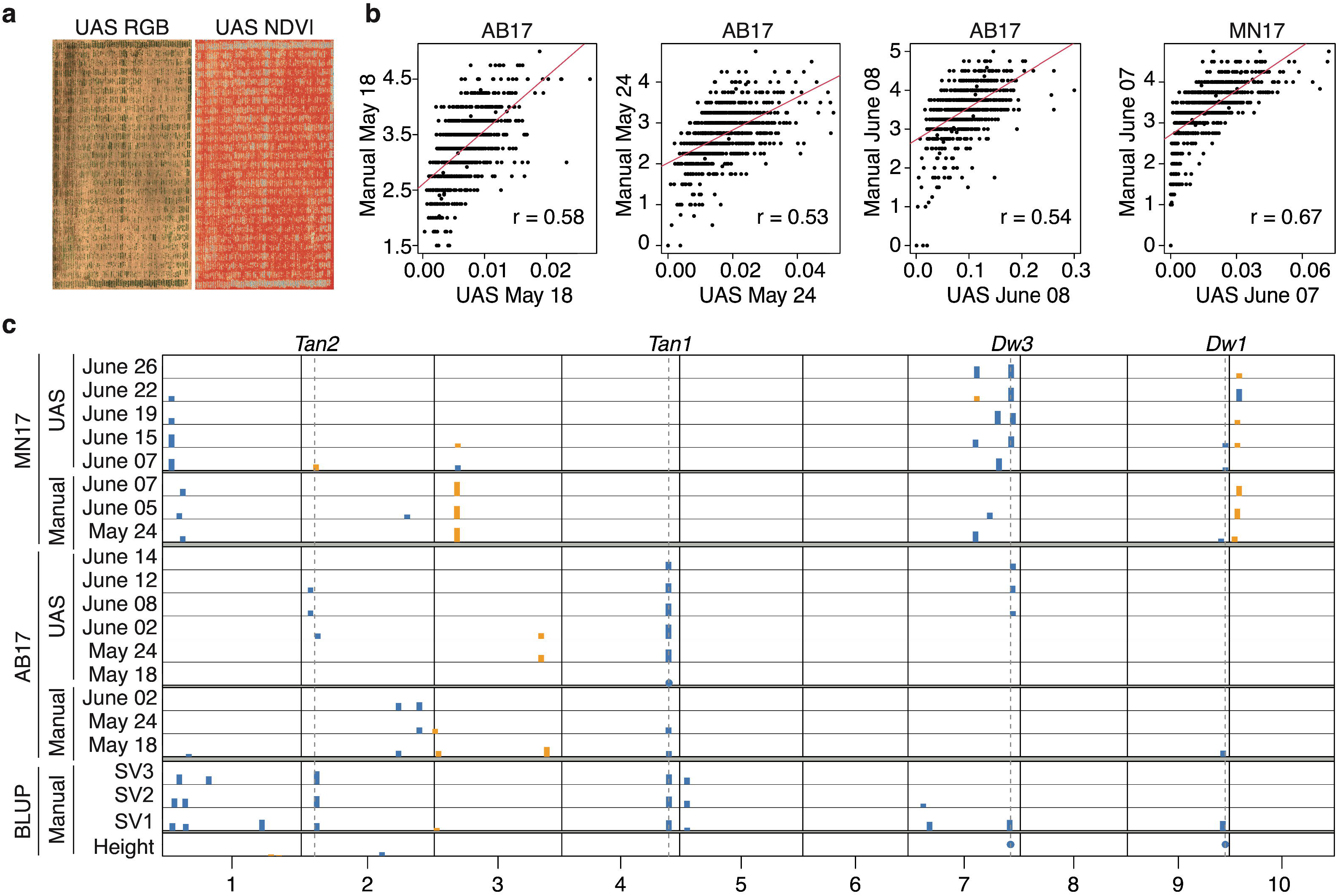
UAS phenomics mapped sorghum chilling tolerance loci that colocalized with classical tannin genes and dwarfing Dw3 gene. **a**) Aerial image of AB17 (Ashland Bottoms 2017) early-planted field trial generated using stitched RGB imagery and NDVI spectrum used for chilling tolerance (CT) phenotyping on June 08, 2017. **b**) Correlation between manual seedling vigor ratings and their corresponding date UAS NDVI values from three AB17 and one MN17 (Manhattan 2017) HTP data. **c**) Joint linkage mapping of UAS NDVI values and manual seedling vigor ratings from MN17 and AB17 field trials. JLM of emergence count and seedling vigor rating BLUPs from six early-planted field trials and plant height BLUPs from one early- and two normal-planted field trials. Positive or negative effects of the BTx623 allele was indicated in orange or blue colors, respectively. The percentage of variation explained is proportional to the height of the rectangular bar for each locus and loci explaining phenotypic variation >10% were noted with circles. Abbreviations: EC, emergence count; SV1–3, seedling vigor1–3.

JLM of NDVI values from AB17 identified four QTL, with each QTL explaining 2.5– 7.5% of the phenotypic variation (Figure 1c). In total, the QTL explained 7–13% variation for the NDVI values (Table S2). The chromosome 4 CT QTL was consistently detected across different NDVI values. The QTL on chromosomes 2 and 7 were mapped with three NDVI values, and the QTL on chromosome 3 was mapped with two NDVI values. Positive alleles were inherited from the Chinese founders, except for the allele on chromosome 3. The QTL on chromosome 4 colocalized (<0.2 Mb) with the classical tannin *Tan1* gene. The QTL on chromosomes 2 and 7 were mapped ~2 Mb and ~4 Mb from the classical tannin *Tan2* and the dwarfing *Dw3* gene, respectively. The four CT QTL detected with AB17 NDVI values mapped to the same genomic regions identified using manual seedling vigor ratings from best linear unbiased predictions (BLUPs) generated using six early-planted field trials. The QTL on chromosome 2 was mapped with NDVI values from a single field season, AB17, while seedling vigor rating BLUPs from six field seasons were required to detect the same QTL.

JLM of MN17 NDVI values mapped eight QTL, with each QTL explaining 2–7% of the phenotypic variation (Figure 1c). These QTL in total explained 18–24% variation for NDVI values in MN17 (Table S3). The positive alleles on chromosomes 1 and 7 were derived from the Chinese founders. Although the MN17 field trial was planted early, the seedlings emerged under optimal conditions and experienced chilling stress one week later. Due to this year-to-year environmental variation, the QTL mapped with MN17 NDVI values were different from AB17 QTL, except for the chromosome 7 QTL mapping close to the *Dw3* gene. The MN17 NDVI QTL detected were consistent with the mapped loci of MN17 manual seedling vigor ratings. The QTL on chromosome 7, close to the *Dw3* gene, was mapped with NDVI values from one month after seedling emergence in both AB17 and MN17 field trials. Overall, the QTL mapped with NDVI values were consistent with the manual seedling vigor rating QTL reported in Marla et al. (2019). Due to fluctuating natural chilling stress between locations and years, the effect size of CT QTL was reduced (Figure 1c). However, these small detectable QTL effects may directly translate into improved chilling fitness in target environments (Cobb et al., 2018).

### Two of the four first-generation markers functioned in diverse genetic backgrounds

Composite interval mapping of seedling vigor ratings from AB16 mapped CT QTL on chromosomes 1, 2, 4, and 9. Four KASP genotyping markers (S1_08641374, S2_08884669, S4_61442862, and S9_56611539), generated using the peak QTL SNPs, were developed for marker-assisted breeding. These markers were effective in differentiating the Chinese founders-derived CT allele from the US-derived alternate BTx623 allele. In the USDA-CSRL sorghum breeding program, the KASP marker on chromosome 2 (S2_08884669) revealed the presence of CS allele in 12 elite US sorghum lines and CT allele in three Chinese lines (Figure 2a). The intercross between BTx642 and (B403 × HKZ) F_1_ progeny [BTx642 × (B403 × HKZ)] generated plants with CS/CS or CS/CT alleles in a 1:1 expected ratio (Figure 2b). Self-pollinating heterozygous plants generated progeny that segregated in a expected ratio of 1: 2: 1 (χ^2^ *p*-value 0.53) (Figure 2c). In the second population, BTx642 / (BTx398*ms3* / (BTx623 / Kaoliang)- Sel) intercross progeny contained individual plants with CS/CS or CS/CT alleles. Self-pollinated heterozygous (CS/CT) plants generated progeny that segregated in a expected ratio of 1: 2: 1 (χ^2^ *p*-value 0.68). Similar to S2_08884669, marker S1_08641374 effectively differentiated the CT allele from the CS allele in two segregating populations generated by selfing two randomly selected individual plants for desirable traits (Figures S1a–c).

**Figure 2:**
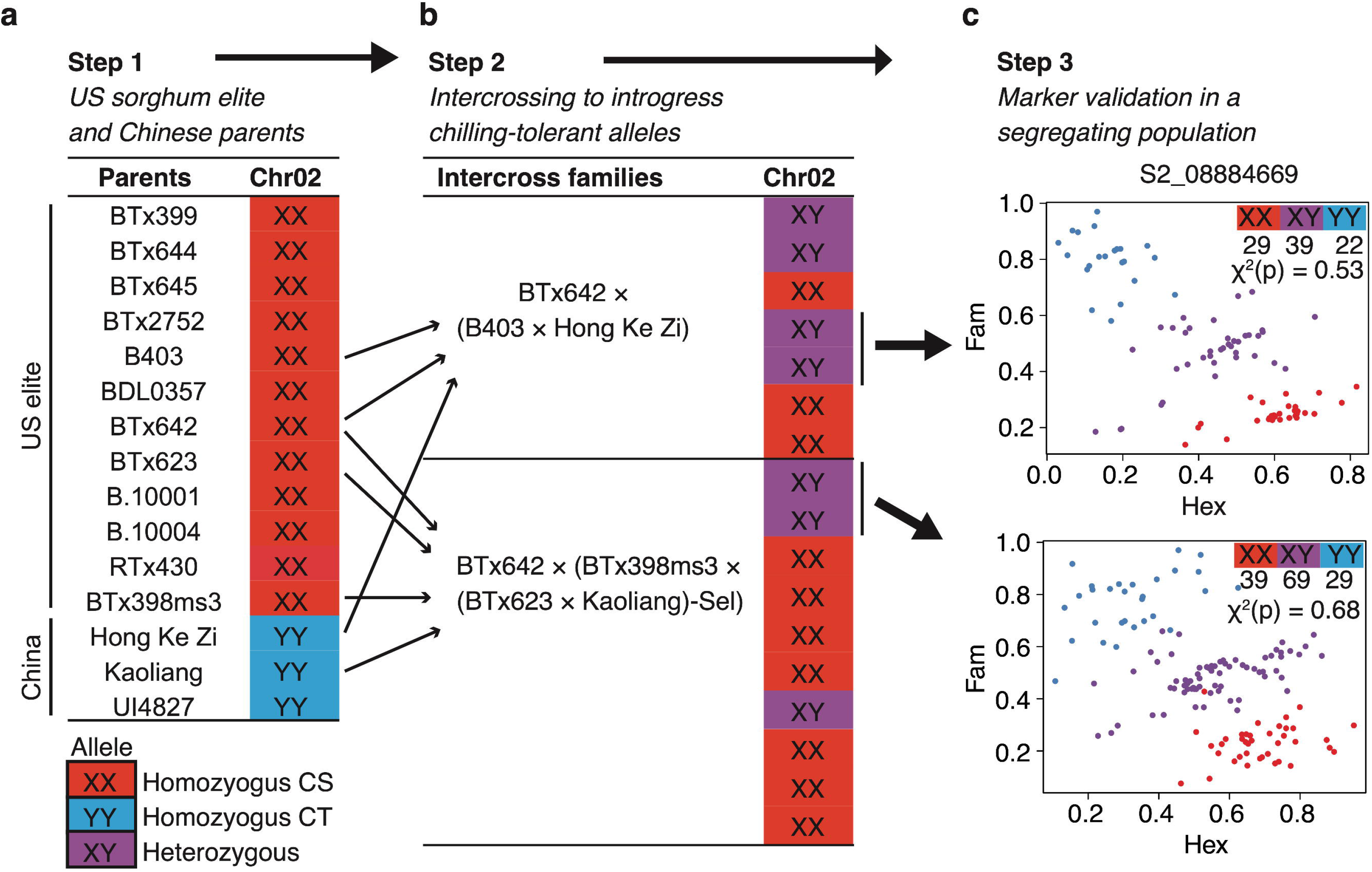
First-generation marker differentiated the US vs. Chinese allele in the USDA-CSRL sorghum breeding program. **a**) Step 1: First-generation marker, designed based on the chilling-tolerant (CT) QTL peak SNP on chromosome 2 (S2_08884669), identified the presence of donor allele in the Chinese parents and the alternate allele in the US elite parents. **b**) Step 2: Intercrosses were conducted between the US and Chinese parents to introgress CT alleles into the US lines. Arrows indicate the parents used for each intercross. Two intercross populations, [BTx642 × (B403 × Hong Ke Zi)] and [BTx642 × (BTx398*ms3* × (BTx623 × Kaoliang)-Sel)] genotyped with S2_08884669 marker showed individual plants carrying either homozygous alternate allele (XX) or heterozygous for the donor and the alternate allele (XY). **c**) Step 3: The progeny of two selfed heterozygous (XY) plants segregated in an expected ratio of 1:2:1 for XX:XY:YY (χ^2^ *p*-values 0.53 and 0.68). Arrows indicate the individual plants that were selfed to generate each intercross population.

By contrast, first-generation markers S4_61442862 and S9_56611539 markers failed to differentiate the US vs. Chinese parents (Figures S1d and 1e). Allelic distribution of the four KASP SNP markers showed missing (N) allelic information in 20–60% of the 30 Chinese and 390 SAP lines (Figure S2 and Figure 3a). The CT allele for SNPs S1_08641374, S2_08884669, and S9_56611539 was the only detected allele in the Chinese lines (Figure S2). In the SAP, CT alleles for S1_08641374, S2_08884669, and S9_56611539 were observed at 12%, 16% and 6% frequency, respectively. In the Chinese lines, the CT allele for S4_61442862 was observed at 68% and the CS allele at 16% frequency. In the SAP, the CT allele S4_61442862 was detected in 20% of the lines. Allelic distribution of S4_61442862 in 1813 georeferenced sorghum lines, a subset of the 10K GBS lines (Hu et al., 2019), revealed the CT allele was a common allele in the global germplasm (Figure 3a). In summary, the first-generation markers designed to introgress CT alleles were designed based on polymorphisms that were common alleles in the global germplasm.

**Figure 3:**
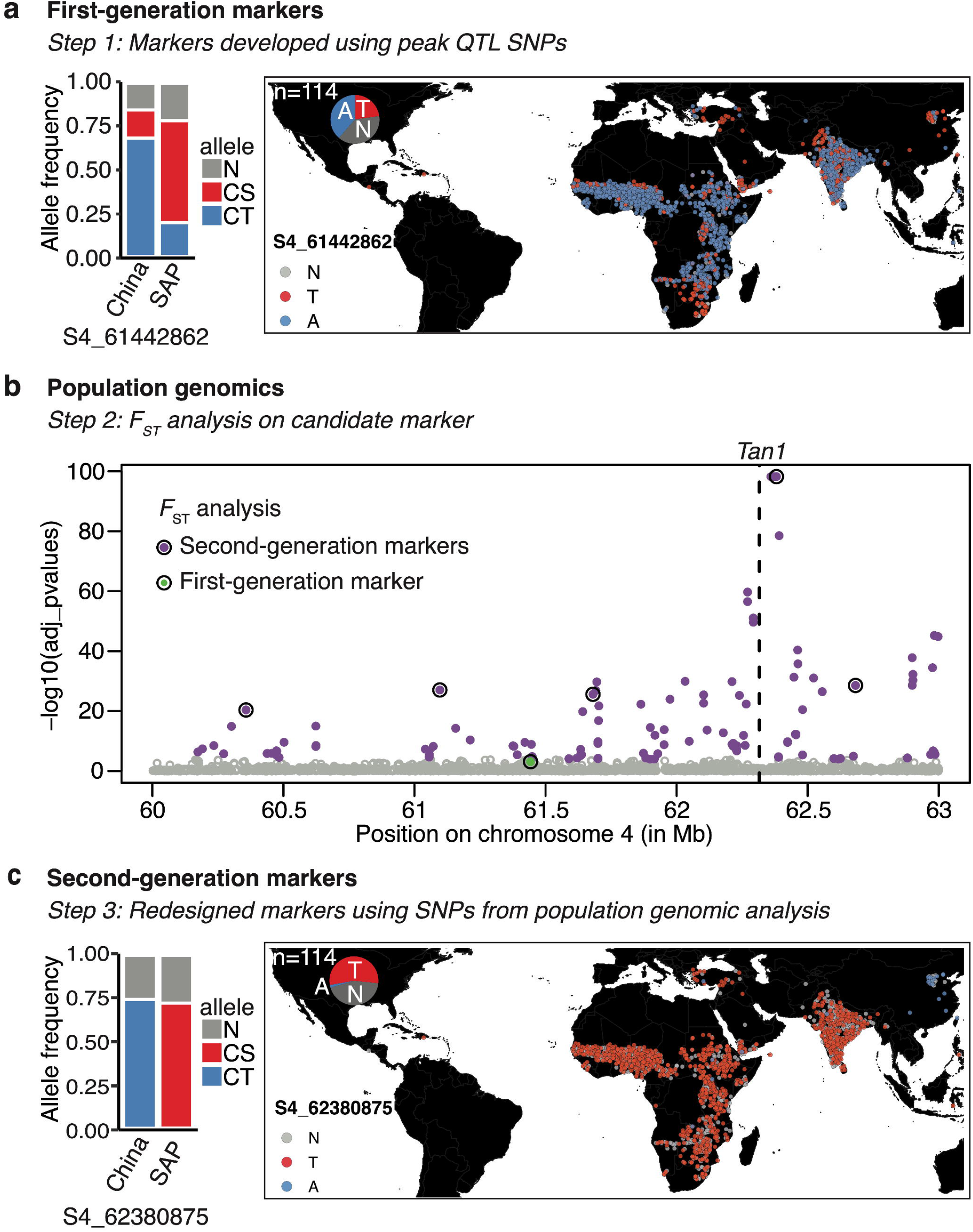
Population genomics enabled detection of polymorphic SNP alleles common in Chinese lines but rare in the global germplasm. **a**) Allele frequency of the first-generation SNP marker S4_61442862 in 30 Chinese and 390 SAP lines. Global allelic distribution of the CT allele (A) *vs*. the alternate allele (T), and missing genotype information (N) in 1813 georeferenced sorghum lines, which did not contain any US lines. As the 114 US lines in the sorghum 10K lines do not have georeferences, they were presented as a pie chart. **b**) *F*_ST_ analysis conducted on 60–63 Mb region of chromosome 4 using the R OutFLANK package. Outlier loci in the selected genomic regions were highlighted in purple, first-generation marker was colored in green, and second-generation markers from the outlier SNPs were highlighted with a circle. The *Tan1* gene at 62.3 Mb on chromosome 4 was noted with a black dashed line. **c**) Allele frequency of the second-generation SNP S4_62380875 in 30 Chinese and 390 SAP lines, and its global allelic distribution in 1813 georeferenced sorghum lines.

### Population genomics analysis enabled identification of second-generation markers

Second-generation KASP markers were designed based on the high-resolution mapping loci detected using JLM analysis with 43K SNPs and 771 RILs (Marla et al., 2019). Given that the first-generation markers were designed using globally common CT alleles, *F*_ST_ test was conducted on SNPs within the loci of interest to identify alleles common in the Chinese germplasm but were rare in the global germplasm. For e.g. *F*_ST_ analysis of 60–63 Mb region on chromosome 4 with 30 Chinese and 390 SAP lines revealed an average *F*_ST_ of 0.11. Based on the Bonferroni-adjusted *p*-value < 0.01, *F*_ST_ analysis identified 128 outlier SNPs in the genomic regions of interest (Figure 3b). The most extreme *F*_ST_ outliers on chromosome 4 colocalized with the *Tan1* gene.

Five KASP markers, one at the QTL peak (S4_62380875) and four flanking markers (S4_60623655, S4_61096729, S4_61680898, and S4_62682585) were developed to introgress the CT allele into chilling-sensitive US sorghums (Table S1). Among these markers, CT allele was the common allele in the Chinese lines while being rare in the SAP (Figure 3c and Figure S3a). Allelic distribution of the peak *F*_ST_ outlier SNP (S4_62380875) in 1813 geo-referenced lines revealed the second-generation marker CT allele was the common allele in Chinese lines but rare in the global germplasm (Figure 3c). Similar to chromosome 4 QTL, *F*_ST_ analysis was used to develop second-generation KASP markers to introgress CT loci mapped on chromosomes 1, 2, and 9 (Figures S3b, S4a, and S4c). Allelic frequency of the markers on chromosomes 1, 2, and 9 revealed CT allele was the common allele in the Chinese lines while being rare in the SAP (Figures S3c, S4b, and S4d).

### Second-generation markers accurately detected the CT allele in diverse USDA-CSRL breeding lines

Second-generation markers, developed using CT alleles that were globally rare but common to locally adapted Chinese germplasm, were effective in differentiating the NSZ × BTx623 RIL CT allele vs. the alternate allele in chilling-sensitive US elite lines from the USDA-CSRL breeding program (Figure 4a). The F_2_ population generated by selfing the progeny derived from the genetic cross between BTx2752 and NSZ RIL, carrying Chr1+, Chr2+, and Chr4+ CT alleles, segregated in an expected ratio of 1: 2: 1 (χ^2^ *p*-value = 0.91) with marker S4_62380875. In a second F_2_ population generated from crossing BTx642 with the NSZ RIL, we observed a segregation ratio of 1: 2: 1 ratio (χ^2^ *p*-value = 0.42) with S4_62380875. Similar to S4_62380875, second-generation markers on chromosomes 1 and 2 (S1_07620913 and S2_07404837) segregated these two F_2_ populations in an expected ratio of 1: 2: 1 ratio (Figure S5). Despite generating chromosome 9 markers, this locus was not used in chilling tolerance breeding due to its colocalization with the undesirable tall plant *Dw3* allele.

**Figure 4.**
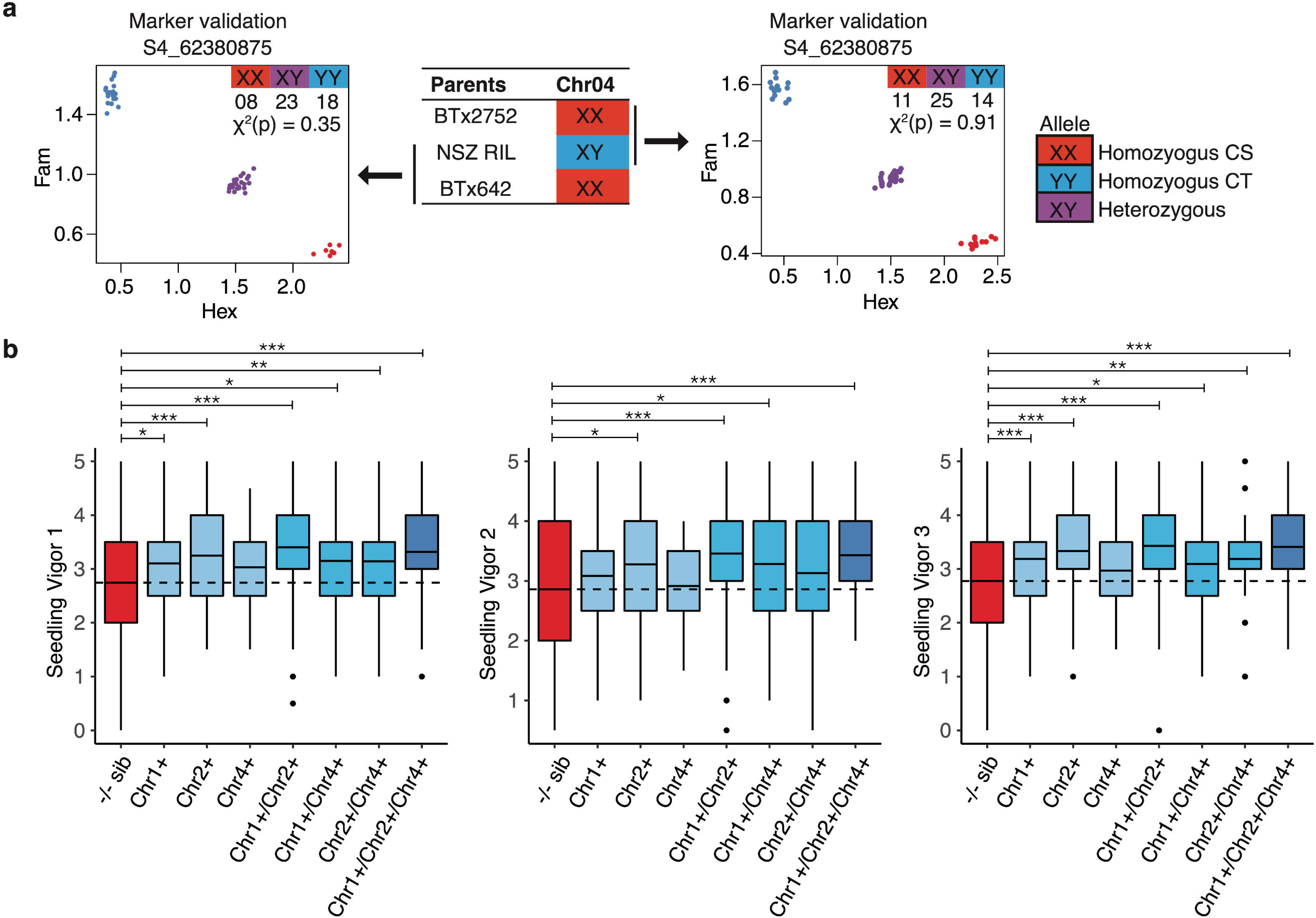
Testing of segregants in independent sorghum breeding programs validated the utility of second-generation markers in improving early-season chilling tolerance. **a**) Second-generation marker S4_62380875, generated using outlier *F*_ST_ SNPs, identified the presence of donor allele in the BTx623 by NSZ RIL and the alternate allele in two US elite lines. Two intercross F_2_ populations, generated by selfing the F_1_ progeny of BTx642 × NSZ RIL and BTx2752 × NSZ RIL, segregated in an expected ratio of 1:2:1 for XX:XY:YY (χ^2^ *p*-values 0.42 and 0.91). **b**) Second-generation markers were used to identify the F_2_ plants with different CT allele combinations from a segregating population generated by crossing three WKARC lines with two chilling NAM RILs. The F_3_ families with different CT alleles were screened for their early-season chilling response under natural chilling stress in Hays 2019. In the F_3_ families, control −/− sib has no CT alleles, single CT allele families carried Chr1+, Chr2+, or Chr4+ from chromosome 1, 2, or 4, two CT alleles stacked families carried Chr1+Chr2+, Chr1+Chr4+, or Chr2+Chr4+ allele combinations, and the three CT alleles stacked family carried Chr1+Chr2+Chr4+. Three seedling vigor ratings (SV1–3) from week-one, -two, and -four after emergence were included. The mean of each F_3_ family was represented as a black horizontal line in the box plots. Lines above box plots indicate pairwise comparisons with a statistically significant differences with *p*-value was less than 0.05 (*), 0.01 (**), and 0.001 (***).

### CT alleles improved early-planted seedling vigor in the WKARC sorghum program

To evaluate if the second-generation markers improved early-season chilling tolerance in an independent breeding program, these markers were used to genotype a segregating F_2_ population derived from a genetic cross between three WKARC breeding lines and two CT NAM RILs. Under natural chilling stress field trials, the F_3_ family with CS alleles at 1, 2, and 4 CT loci (No CT allele, −/− sib) contained lower average seedling vigor (SV1–3) ratings compared to the F_3_ families with CT alleles (Figure 4b). The F_3_ family with chromosome 2 CT loci (Chr2+) showed 14–20% increase in average SV1–3 ratings compared to the −/− sib control. The F_3_ family with chromosome 1 CT loci (Chr1+) showed 13% and 15% increased SV1 and SV3 ratings, respectively, compared to the −/− sib control. The F_3_ family with chromosome 4 CT allele (Chr4+) showed no significant differences (*p*-value > 0.05) in SV1–3 ratings compared to the −/− sib family. The F_3_ family carrying two CT alleles (Chr1+Chr2+ or Chr1+Chr4+) showed a significant 12–24% increase in SV1–3 ratings compared to −/− sib control, while the family with Chr2+ Chr4+ showed 14% improved SV1 and SV3 ratings. Stacking all three CT alleles (Chr1+Chr2+Chr4+) together provided 19–23% higher SV1–3 ratings, compared to the −/− sib control. Taken together, these results demonstrated a significant (*p*-value < 0.05) increase in SV ratings, under natural chilling stress, in the F_3_ families with CT alleles at chromosome 1, 2, and 4 CT loci.

### Marker-assisted breeding increased early-planted SV in US sorghum hybrids

In early-planted field trials, sorghum hybrids developed by crossing inbred NILs, with CT alleles at chr2+, chr4+, or −/− sib, with five nuclear *male-sterile3* (*ms3*) elite parents contained similar agronomic characteristics as the US elite inbreds at maturity (Figure 5a). As expected, early-planted seedling performance traits (EC, SV1, and SV2) were significantly higher (40–50%, *p*-value < 0.001) in the Chinese parents compared to the US elite parents (Figure 5b and Figure S6), validating the phenotyping experiments. Among the inbred NILs, average EC, SV1 and SV2 ratings were higher in inbreds with Chr2+ or Chr4+ compared to −/− sib. However no significant statistical differences (*p*-value > 0.05) were observed between inbred NILs. In the hybrids, no significant difference was observed with EC between −/− sib, chr2+, or chr4+ hybrids (Figure 5b). Similarly, no significant differences were observed with SV1 in the hybrids (Figure S6). Significant increase in SV2 (14%, *p*-value = 0.04) was observed only between hybrids carrying chr4+ and the −/− sib, while no significant difference was present between chr2+ and −/− sib hybrids (Figure 5b).

**Figure 5.**
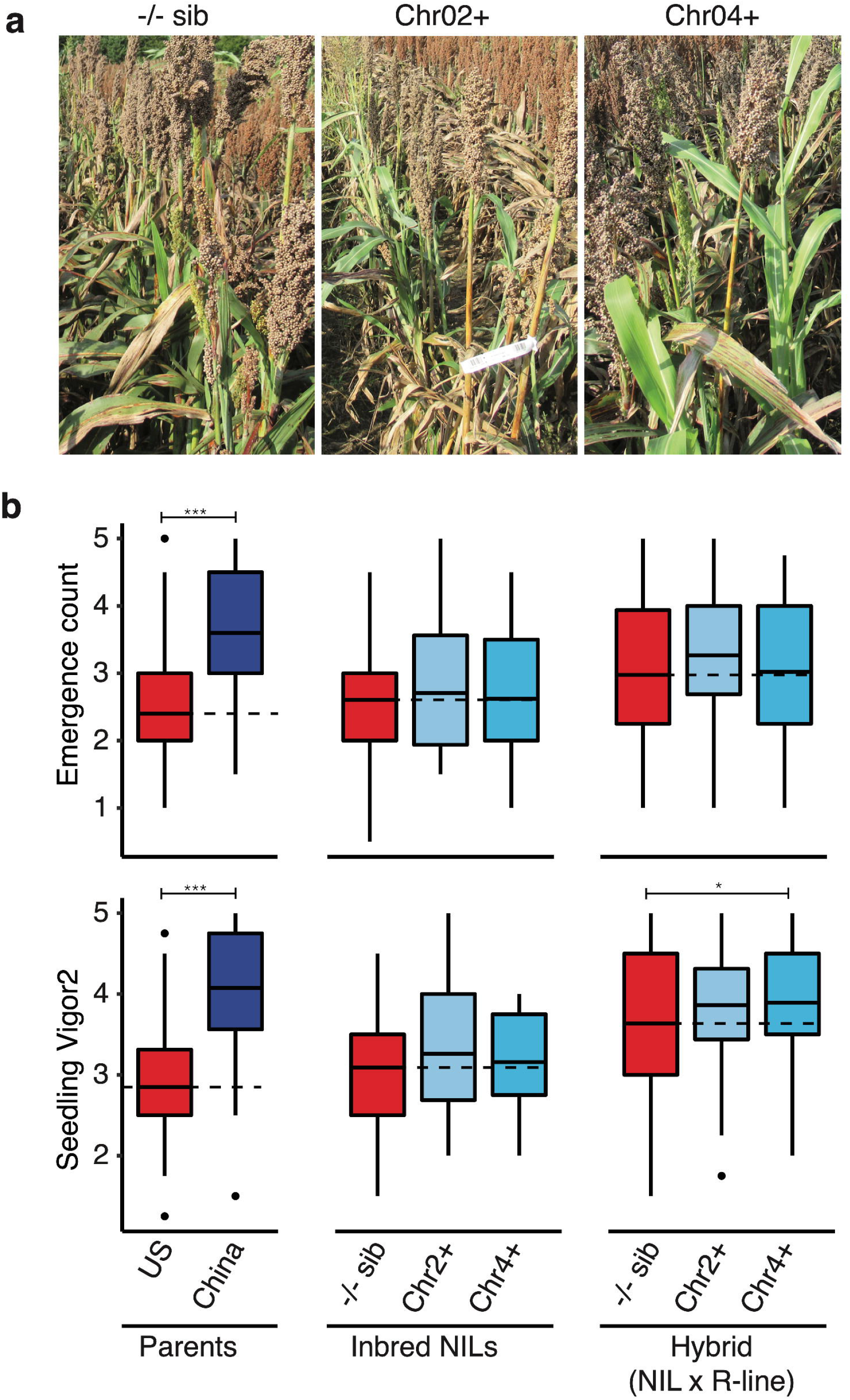
Marker-assisted breeding for chilling tolerance increased early-planted seedling performance trait ratings in sorghum hybrids. **a**) Representative field images of sorghum hybrids with Chr2+ or Chr4+ CT alleles and their control −/− sib not carrying any CT alleles. **b**) Comparison of CT alleles performance in inbred near-isogenic lines (NILs) and hybrids carrying no CT allele (−/− sib), Chr2+, or Chr4+ CT allele. Hybrids were generated by crossing five US elite R lines carrying the *male sterility3* (*ms3*) gene with inbred NILs. The US and Chinese parents were included as controls. Emergence count and seedling vigor2 (SV2) showed an increase in inbred NILs and hybrids with CT QTL compared to −/− sib. In hybrids, only SV2 contained a significant increase compared to the −/− sib control.

## DISCUSSION

### UAS phenomics-enabled prebreeding for a complex adaptive trait

Breeding for abiotic stress tolerance was slowed down by the complex genetic architecture of stress tolerance, for e.g. sorghum chilling tolerance (Figure 1c), and lack of uniform stress under target environments between years and locations (Collins et al., 2008; Vandenbroucke & Metzlaff, 2013). Additionally, complex trait breeding was impeded by poor scalability due to the time-consuming and tedious nature of manual phenotyping (Araus et al., 2018). For example, improving scalability for sorghum CT was limited by the amount of time (~5 h) required for manually phenotyping each seedling performance trait in a field trial. In this study, UAS phenotyping required 45 min of UAS image capture and NDVI values were extracted in ~2 h for each trait. JLM with UAS NDVI values mapped CT loci that colocalized with manual seedling vigor CT loci (Figure 1c), validating the utility of UAS phenomics in mapping the genetic architecture of complex adaptation traits. UAS phenomics from AB17 field season mapped chromosome 2 QTL while BLUPs from six field seasons were required to consistently map the same QTL (Figure 1c), suggesting increased power of UAS phenomics in complex trait mapping.

Given the ease of UAS image acquisition and mapping common CT QTL with UAS and manual phenotyping, improved scalability with UAS phenomics could lead to strong QTL-to-marker associations in applied breeding programs for developing climate-resilient crops (Varshney et al., 2021). Application of UAS phenomics was limited to upstream genetic studies for high heritability traits, such as plant height, lodging, and disease resistance (Wang et al., 2018; Singh et al., 2019; Sarkar et al., 2020; Zhou et al., 2020; Zhang et al., 2021). Our study directly demonstrates the utility of UAS phenomics for prebreeding of complex traits.

### Population genomics generated markers for improved trait predictability

Advances in genomics and quantitative genetics enabled identification of thousands of QTL in public breeding programs, however, most have not yet been used in molecular breeding (Bernardo, 2016; Mace et al., 2019). Weak trait-to-marker associations, due to low-resolution mapping with bi-parental families, low marker density, and complex LD in the public breeding programs, limited the utilization of mapped QTL in developing improved varieties (Cobb et al., 2018). In this study, utilizing a chilling NAM population with 43K GBS SNPs (Marla et al., 2019) and HTP provided high-resolution CT mapping (Figure 1c) addressed weak trait-to-QTL association from previous sorghum chilling tolerance studies (Knoll et al., 2007; Burow et al., 2010). For successful implementation of a public marker development program, the markers developed should function across elite breeding lines from different breeding programs (Bernardo, 2016; Cobb et al., 2018). Two of the four first-generation markers failed to differentiate the Chinese vs. US elite lines in the USDA-CSRL program (Figure S1b) as the CT allele was commonly present in diverse sorghum lines (Figure 3a), indicating the need to develop markers with improved QTL-to-marker association to accurately identify the target allele across diverse genetic backgrounds.

Leveraging population genomic approaches to identify the genomic regions selected in a locally-adapted germplasm and utilizing the polymorphic alleles common in local germplasm but rare globally has the potential to improve marker functionality in diverse breeding programs (Muleta et al., 2019). Second-generation KASP markers, developed based on *F*_ST_ analysis outlier SNPs, differentiated the Chinese vs. US elite lines in the USDA-CSRL program (Figures 3c, 4a, and Figure S5), indicating the potential of these markers to function in independent public sorghum breeding programs. Although first-generation markers were designed using single-family mapping QTL SNPs and without prior *F*_ST_ analysis, two of the four markers differentiated the Chinese vs. US elite lines in the USDA-CSRL program because the two SNPs used for marker development were *F*_ST_ outliers (Figures S3 and S4).

Overall, integrating population genomic analysis in marker development for sorghum CT improved QTL-to-marker association, critical for markers to function in different public marker-assisted breeding programs (Cobb et al., 2018). Demonstrating strong marker-to-trait association with second-generation markers, marker-assisted selection improved early-planting SV ratings in the WKARC sorghum breeding program and SV2 in sorghum hybrids with Chr4+ allele in the KSU sorghum breeding program (Figures 4b and 5b). Taken together, increased seedling chilling tolerance in early-planted field trials showed population genomics-assisted markers developed as a proxy for chilling tolerance improving trait predictability.

### Combining genomics and phenomics for complex adaptive trait breeding

Despite profound knowledge on the genetic architecture of complex traits, limited success stories exist on molecular breeding for complex traits due to the failure of genetics under controlled conditions translating to target population of environments (TPE), weak trait-to-marker associations, and the functioning of markers and QTL in diverse breeding programs (Blum, 2014; Cobb et al., 2018; Simmons et al., 2021; Ruiz-Vera et al., 2022). This sorghum chilling tolerance presents a case study demonstrating the utilization of genome-to-phenome approaches for complex trait improvement (Figures 6a and 6b). Conducting sorghum CT research in natural chilling stress field trials (Figures 1a and 6b), to replicate relevant spatial and temporal stress variation endured in target environments, addressed the possible failure of controlled conditions genetics translating to TPE in previous studies (Simmons et al., 2021; Ruiz-Vera et al., 2022).

**Figure 6.**
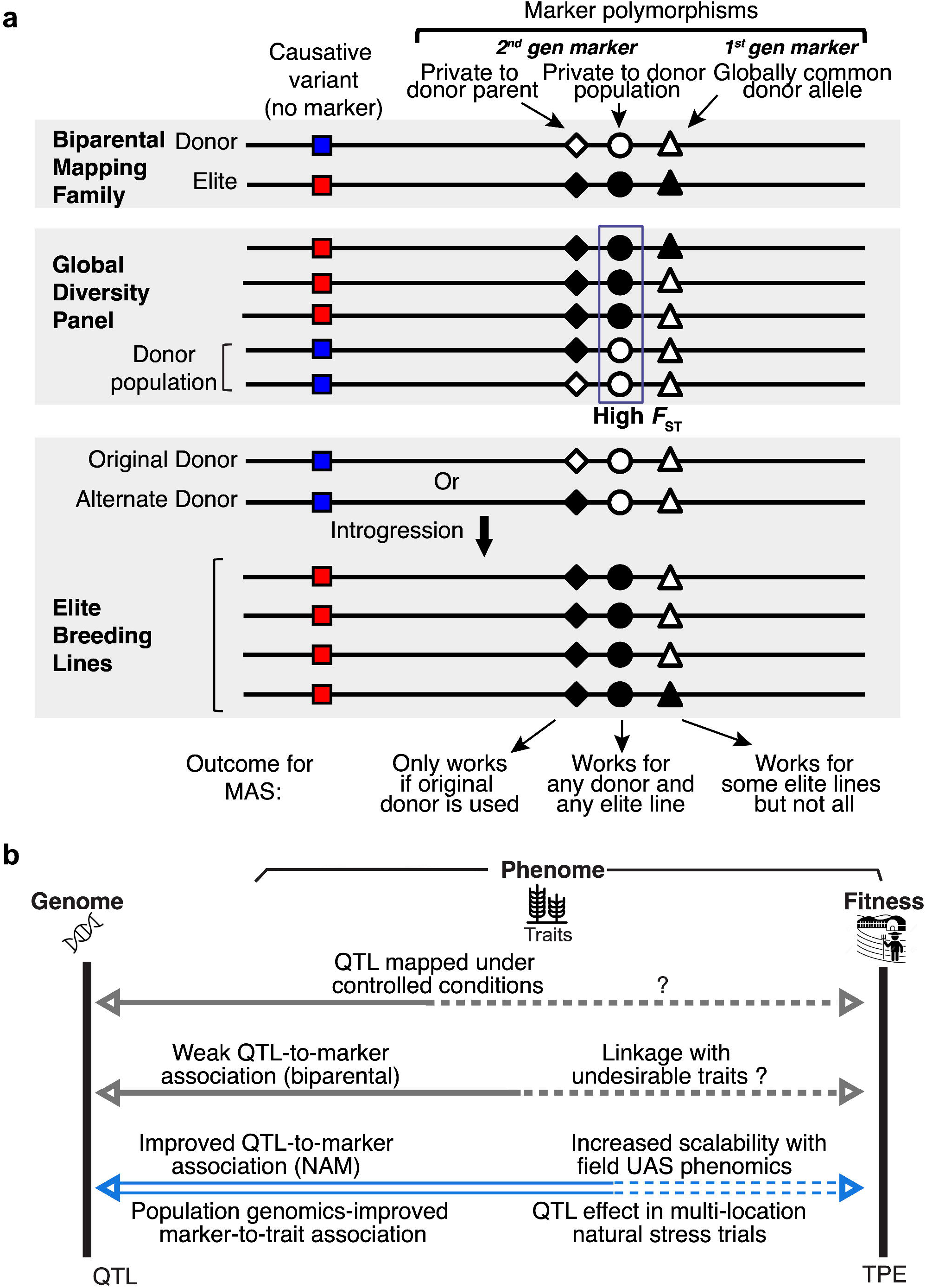
Integrating genomics and phenomics for prebreeding of complex adaptive traits. **a**) Schematic representation of marker-assisted breeding for introgressing CT alleles into diverse breeding programs. Top panel depicts the allelic distribution of three markers, that are in linkage disequilibrium with the causative variant, in a biparental mapping family. Panels two and three represent the allelic distribution of markers in a global diversity panel and elite breeding lines, respectively. All three markers will function effectively with the mapping population in which the QTL was detected, however, functionality of these markers in diverse breeding programs depends on the SNP polymorphism commonality in global germplasm. First-generation markers function with the biparental mapping family but fail in different breeding programs with diverse lines as the CT allele is common in global germplasm. Molecular markers generated to identify a private parent allele function only when the same donor parent is used for trait introgression into different breeding programs. Second-generation markers, designed on population genomics outlier SNP alleles conserved in the donor population but rare globally, can function in global diversity panels and elite breeding lines. Second-generation markers private to the donor population can be effective in introgressing desirable alleles into different breeding programs. **b**) Genome-to-phenome model depicting for complex adaptive trait improvement. Previous breeding efforts for complex trait failed due to failure of QTL mapped in controlled conditions failing to function under target population of environments (TPE) and weak QTL-to-marker associations due to biparental family low-resolution mapping (indicated in gray arrows). Integrating new genome-to-phenome approaches, that provide improved scalability, QTL-to-marker association, and marker-to-trait association, and the introgressed alleles functioning under TPE in marker-assisted breeding, for marker-assisted breeding of complex adaptive traits under TPE could lead to development of improved climate-resilient crops.

High-resolution mapping, obtained with the chilling NAM population and high marker density, resolved the earlier problem of weak trait-to-marker associations (Burow et al., 2010; Bouchet et al., 2017; Marla et al., 2019). Markers developed using the donor parent private allele can differentiate the rare vs. alternate alleles in a breeding program (Figure 6a), however, rare allele-specific markers only function when the same private donor parent was used for trait improvement across breeding programs (Cobb et al., 2018). Second-generation markers developed using private alleles to the donor population (Figures 3c and 6a) were effective in detecting polymorphisms in the two independent sorghum programs (Figure 4a and Figure S5), suggesting these markers can be used for marker-assisted breeding in different public CT sorghum breeding programs.

Despite marker-assisted breeding precisely introgressing target loci into elite lines, breeding efforts for complex trait improvement may not be successful due to the failure of introgressed QTL to provide the desired trait in new genetic backgrounds (Cobb et al., 2018). Improved seedling CT traits in the WKARC breeding program demonstrated the introgressed CT allele to function in a different genetic background than the mapping population (Figure 4b). Additionally, improved SV2 rating in the testcross hybrids with Chr4+ CT allele (Figure 5b), compared to the −/− sib hybrid, indicating that the CT allele functions in sorghum hybrids, commonly used by private commercial US grain sorghum companies (Pfeiffer et al., 2019). Overall, leveraging G2P approaches for sorghum early-season CT addressed previous limitations of low-resolution mapping of complex trait genetic architecture, weak trait-to-marker associations, and markers failing to function in independent public breeding programs (Cobb et al., 2018; Varshney et al., 2021). Our findings from sorghum CT research and development indicate G2P approaches have the potential to successfully work with other complex adaptation traits in different crops (Bernardo, 2008; Bohra et al., 2020).

## EXPERIMENTAL PROCEDURES

### Chilling NAM population

The chilling NAM population of 771 RILs was generated by crossing the US sorghum reference line BTx623 with three chilling-tolerant Chinese founders, Niu Sheng Zui (NSZ; PI 568016), Hong Ke Zi (HKZ; PI 567946), and Kaoliang (Kao; PI 562744) (Marla et al., 2019). The chilling NAM population development, genotyping-by-sequencing (GBS), and single nucleotide polymorphism (SNP) variant calling was explained in more detail in Marla et al., (2019). The chilling NAM GBS data combined with previously published *Ape*KI GBS data from ~10,000 diverse lines (Hu et al., 2019) was aligned to the BTx623 reference genome v3.1 (McCormick et al., 2018). SNP calling using Tassel 5.0 GBS v2 pipeline (Glaubitz et al., 2014) retained 43,320 SNPs and 750 RILs for joint linkage mapping. Selected chilling NAM RILs (15FS005_NSZ, 14FS205_Kao, 14FS125_Kao, 15FS083_NSZ, 15FS032_NSZ, 15FS679_HKZ, 15FS698_HKZ, and 14FS273_Kao) with desirable agronomics and different combinations of CT alleles were used in marker-assisted chilling tolerance pre-breeding.

### Early-planted field trials

The chilling NAM population was planted under natural chilling stress in six early- and two normal-planted field trials in 2016, 2017, and 2018 in Kansas, as described in Marla et al., (2019). Each field trial contained two replicates of the NAM population. Early-planted field trials were sown in April or early May, 30–45 days earlier than normal sorghum planting in Kansas (Grain Sorghum Production Handbook 1998). Manual phenotyping for seedling vigor rating was conducted in all eight field trials. UAV phenotyping was conducted in two field trials, AB17 (Ashland Bottoms 2017, 39.14N −96.63W) and MN17 (Manhattan 2017, 39.21N −96.60W).

Six F_3_ families, generated by crossing two chilling NAM RILs with three lines from the Western Kansas Agricultural Research Center (WKARC, Hays, Kansas) sorghum breeding program, were screened for their early-season chilling response in HA19 (Hays 2019, 38.84N, −99.34W). Five replicates of each WKARC F_3_ family, completely randomized within each replication block, were early-planted on April 17. CT hybrids, from the Kansas State University (KSU) sorghum genetics and molecular breeding program (Manhattan, Kansas), were early-planted in 2019 at two locations: Hays and Manhattan. Each field trial consisted of three replicates of the parents, inbred NILs, and hybrids with Chr2+ or Chr4+ CT allele or without CT allele (−/− sib). CT hybrid evaluations were early-planted on April 19 in HA19 and April 17 in MN19 field trials.

### Manual field phenotyping

In natural chilling stress field trials, seedling performance traits, emergence count (EC) and three seedling vigor (SV) ratings (SV1–3), were scored on a 1–5 scale (1 for low and 5 for high) as described previously (Marla et al., 2019). The SV ratings scale previously described (Maiti et al., 1981; Knoll et al., 2007; Burow et al., 2010) was modified (1 for high and 5 for low SV) for consistency with EC rating. One-week after emergence, EC was determined by counting the number of seedlings that emerged and converted to a scale of 1–5 representing 20, 40, 60, 80, and 100% emergence, respectively. Three SV1–3 ratings, scored independently of emergence count, were collected at week-1, -2, and -4, respectively, after emergence. SV ratings were scored on a rating scale of 1–5 with a rating of 1 for low and 5 for robust vigor. In the HA19 field trial, SV1–3 ratings were collected at week-1, -2, and -4 after emergence. In MN19 and AB19 field trials, EC and SV1 ratings were collected at week-1 after emergence, and SV2 ratings at week-2 after emergence.

### High-throughput phenotyping with UAS and data processing

UAS high-throughput phenotyping (HTP) was conducted on AB17 and MN17 early-planted field trials. HTP data in the 2017 chilling NAM trials were collected by a DJI Matrice 100 quadcopter (DJI, Shenzhen, China) carrying a MicaSense RedEdge-M multispectral camera (MicaSense Inc., USA). HTP data were collected throughout the entire growth cycle but only early-stage data, upto 45 days after emergence, were analyzed in this study. Flight plans were created using CSIRO mission planner application and missions were executed using the Litchi Mobile App (VC Technology Ltd., UK https://uavmissionplanner.netlify.app/). Aerial image overlap rate between two geospatially adjacent images was set to 80% both sequentially and laterally to ensure optimal orthomosaic photo stitching quality. All UAS flights were set at 20 m above ground level at 2 m/s and conducted within 2 h of solar noon. To improve the geospatial accuracy of orthomosaic images, white square tiles with a dimension of 0.30 m × 0.30 m were used as ground control points and were placed at four corners of the field before image acquisition and surveyed to centimeter-level resolution using the Emlid REACH RS+ Real-Time Kinematic Global Navigation Satellite System unit (Emlid Ltd, Hong Kong, China). A semiautomated image processing pipeline developed by Wang et al., (2020) was used to generate field orthomosaic photos and extract plot-level NDVI (Rouse et al., 1974) and crop coverage.

### Statistical analyses

Correlation between AB17/MN17 seedling vigor ratings and UAS NDVI data was determined using averaged values of two replicates from each field trial. Pearson pairwise correlation analysis was performed using the *cor* function in R package. Chi-square test (χ^2^), using R *chisq.test*, was conducted to determine whether there was a significant difference between observed and expected frequencies of CT vs. chilling-susceptible (CS) alleles from KASP genotyping in intercross populations generated from the Chinese and US elite parents. In the WKARC breeding program F_3_ families, statistical comparisons were conducted between the control F_3_ family without any CT alleles (−/− sib) and individual F_3_ families with different CT allele combinations using a Dunnett’s test. Boxplots were generated using ggplot2 R package (Wickham, 2016). To determine if chromosome 2 or 4 (Chr2+ or Chr4+) CT alleles improved seedling performance traits, EC, SV1, and SV2, in CT hybrids, statistical comparisons were conducted between the US and Chinese parents, NILs with Chr2+ or Chr4+ or −/− sib, and hybrids with Chr2+ or Chr4+ or −/− sib using a Dunnett’s test.

### Joint linkage mapping

GBS of the chilling NAM population generated 43K SNPs for JLM analysis. JLM was performed individually for each location with the averaged data of two replicates. JLM of AB17 and MN17 NDVI values was conducted, as previously described (Marla et al., 2019), using the stepwise regression approach in TASSEL 5.0 (Glaubitz et al., 2014). Entry and exit limit of forward and backward stepwise regressions was 0.001 and threshold cut off was set based on 1000 permutations. Allelic effects of individual QTL were expressed relative to the BTx623 allele, where alleles with positive- or negative-additive effects were derived from BTx623 or Chinese founders, respectively.

### Population genomics analysis

A collection of 30 Chinese and 390 sorghum association panel (SAP) lines, a subset of the 10K GBS sorghum lines (Hu et al., 2019), were used to calculate the fixation index for SNPs within the loci of interest. GBS data was extracted from chromosomes 1 (4–12 Mb), 2 (7–11 Mb), 4 (60–63 Mb), and 9 (56.4–57.2) using VCFtools package (Danecek et al., 2011). Imputation was conducted on each genomic region separately using Beagle 4.1 (Browning & Browning, 2016). Pairwise SNP differentiation (*F*_ST_) between the Chinese and SAP lines was calculated, and the outliers loci in the selected genomic regions were detected on the basis of an inferred distribution of neutral *F*_ST_ using the OutFLANK R package (Whitlock et al., 2015). Allelic frequency of the first and second-generation KASP markers in the 30 Chinese and 390 SAP lines was calculated on unimputed GBS data from Hu et al., (2019). A global set of 1813 geo-referenced lines, a subset of geo-referenced GBS lines (Lasky et al., 2015) and 10K GBS sorghum lines (Hu et al., 2019), was used to determine the allelic distribution of first- and second-generation markers. Among the 10K GBS sorghum lines, 114 US lines were used for determining the allelic distribution. World map with allelic distribution of the geo-referenced lines was generated using ggplot2 (Wickham, 2016).

### KASP genotyping

Two six mm diameter leaf punches were collected from three-week-old seedlings for KASP genotyping. Leaf tissue samples were dried using silica-gel beads and shipped to Intertek, Sweden, for genotyping. SNPs for the first-generation markers were selected within the QTL mapped using the AB16 manual phenotyping data. Second-generation KASP markers were designed based on the SNPs identified using JLM, from 2016 and 2017 manual phenotyping data, and SNP outliers detected using *F*_ST_ analysis that were in LD with the peak JLM SNPs. KASP markers were designed using Primer3Plus (Untergasser et al., 2007) based on the sorghum reference genome v3.1 with settings product size <120 bp, annealing temperature between 55°C and 62°C with optimum at 57°C, and paired primers annealing temperature within 1°C of one another (Table S1). Tails for use with the KASP genotyping system (Semagn et al., 2014) were added post primer development, with the HEX-fluorescence designated tail added to the CT allele. DNA extraction and KASP genotyping was conducted at Intertek.

### Chilling tolerance breeding populations

KASP (LGC Biosearch Technologies, Middlesex, UK) genotyping markers were tested on breeding populations from two independent US public sorghum breeding programs, the USDAARS Cropping Systems Research Laboratory (CSRL) (Lubbock, Texas) and WKARC sorghum breeding programs. In the USDA-CSRL program, first-generation KASP markers were tested on intercross breeding populations derived from the genetic crosses between 12 US elite grain sorghum lines (BTx399, BTx644, BTx645, BTx2752, B403, BDL0357, BTx642, BTx623, B. 10001, B. 10004, RTx430, and BTx398*ms3*) and three Chinese lines (Hong Ke Zi, Kaoliang, and UI4827 [PI 408816]). Plastic bag emasculations were conducted on the US elite lines to use them as female parents for generating intercross breeding populations. First-generation markers KASP genotyping results from two breeding populations, [BTx642 × (B403 × Hong Ke Zi)], and [BTx642 × (BTx398*ms3* × (BTx623 × Kaoliang)-Sel)], were included in the manuscript. Second-generation markers were evaluated on F_2_ plants generated by selfing F_1_ individuals derived from the genetic cross between BTx2752/BTx642 and a NSZ RIL carrying CT alleles on chromosome 1, 2, and 4.

In the WKARC sorghum breeding program, second-generation KASP markers were tested on six populations generated by crossing three breeding lines, ARCH 10747-1, ARCH 10747-2, and ARCH 12045, with two chilling NAM RILs (14FS205_Kao and 15FS005_NSZ). The ARCH 10747-2 parental line carried both functional tannin genes, while ARCH 10747-1 and ARCH 12045 carried non-functional *Tan1* and functional *Tan2*. The chilling NAM 15FS005_NSZ RIL carried CT alleles on chromosomes 1, 2, and 4. RIL 14FS205_Kao carried CT alleles only on chromosomes 1 and 2, a recombination event before the chromosome 4 CT QTL resulted in the loss of this CT allele in this line. Due to this recombination, the Chr4+ allele in two populations, 14FS205_Kao × ARCH12045 and 14FS205_Kao × ARCH10747-1, was replaced with no CT allele. KASP-genotyped F_2_ plants were self-pollinated to derive 204 F_3_ families of different CT allele combinations (−/− sib, Chr1+, Chr2+, Chr4+, Chr1+/Chr2+, Chr1+/Chr4+, Chr2+/Chr4+, or Chr1+/Chr2+/Chr4+).

Second-generation KASP markers were further tested on the CT hybrids generated from the KSU sorghum program. Four chilling NAM RILs (14FS125_Kao, 15FS083_NSZ, 15FS679_HKZ, and 14FS273_Kao), carrying different combinations of CT alleles on chromosomes 2 and 4, were backcrossed onto plastic bag emasculated female plants of BTx623 for three generations, and then selfed for one generation (BC2F2) to obtain near-isogenic lines (NILs) with or without CT alleles. Individuals fixed for contrasting CT alleles were identified in the segregating population via KASP markers. CT hybrids were generated by cross-pollinating the BC2F2 NILs onto five nuclear *male sterile3* (*ms3*) US elite lines (RTx2737*ms3*, RTx430*ms3*, RTx436*ms3*, RTx437*ms3*, and SC110*ms3*). Chinese lines Gai Gaoliang (PI 610727), HKZ, NSZ, Kao, and Shan Qui Red (PI 656025) were the chilling-tolerant controls, while BTx623 was the chilling-sensitive control in all early-planted field trials.

## ACKNOWLEDGEMENTS

This research was supported by funding from the Kansas Grain Sorghum Commission (KGSC), Foundation for Food and Agriculture Research (FFAR), and TERRA-REF: US Department of Energy’s Advanced Research Projects Agency-Energy (ARPA-E). The study was carried out using the Beocat high-performance computing facility. This study is a contribution --- from the Kansas Agricultural Experiment Station. We would like to thank Matt Davis and Troy Ostmeyer for excellent technical support. The authors state that they have no conflict of interest to declare for this research.

## SUPPORTING FIGURES

Supporting material available at Figshare: https://doi.org/10.6084/m9.figshare.21358191.v1

*Figure S1: First-generation marker genotyping of the US and Chinese lines in the USDA Lubbock sorghum breeding program*.

**a**) Step 1: First-generation marker, designed based on the chilling-tolerant (CT) QTL peak SNP on chromosome 1 (S1_08641374), identified the presence of donor allele in the Chinese lines and the alternate allele in the US germplasm. **b**) Step 2: Intercrosses were conducted between the US and Chinese parents to introgress CT allele into the US lines. Two intercross populations, [BTx642 × (B403 × Hong Ke Zi)] and [BTx642 × (BTx398*ms3* × (BTx623 × Kaoliang)-Sel)] genotyped with S1_08641374 marker showed individual plants carrying either homozygous alternate allele (XX) or heterozygous for the donor and the alternate allele (XY). **c**) Step 3: The segregation pattern of CT/CS alleles observed in the progeny from two randomly selected plants suggest the selfed plants contained XX and XY alleles. **d**) First-generation markers on chromosome 4 and 9 failed to differentiate the US and Chinese parents as the CT allele is common in the global germplasm. **e**) The two intercross populations did not segregate for the marker on chromosome 9 as this population was selected for the *dw1* allele conferring short plant stature.

*Figure S2: Allele frequency of first-generation markers showing the donor allele was common in global diversity*.

Allele frequency of three first-generation markers S1_08641374, S2_08884669, and S9_56611539 was calculated for 30 Chinese and 390 sorghum association panel (SAP) lines separately. Chilling tolerant allele was the only allele identified in the Chinese lines. The presence of CT allele in the SAP indicated the CT allele was not a globally rare allele.

*Figure S3: Population genomics identified markers targeting chilling-tolerant alleles common in locally-adapted lines but rare in global germplasm*.

**a**) Allele frequency of four second-generation markers on chromosome 4, calculated for 30 Chinese and 390 sorghum association panel (SAP) lines separately, showed CT allele was a globally rare allele but a common allele in Chinese lines. **b**) *F*_ST_ analysis conducted on 4–13 Mb region of chromosome 1 using R OutFLANK package. Outlier loci in the selected genomic regions were colored in purple, the first-generation marker in green and highlighted with a circle, and second-generation KASP markers were highlighted with a circle. **c**) Allele frequency of five second-generation markers in 30 Chinese and 390 SAP lines.

*Figure S4: Population genomics-enabled markers identified chilling-tolerant alleles fixed in locally-adapted* lines *but were globally rare*.

**a**) *F*_ST_ analysis of 7–11 Mb on chromosome 2, outlier loci in this region were colored in purple, the first-generation marker indicated in green and highlighted with a circle, and second-generation markers were highlighted with a circle. *Tan2* gene at 7.9 Mb was noted with a black dashed line. **b**) Allele frequency of four second-generation markers on chromosome 2 in 30 Chinese and 390 SAP lines. **c**) *F*_ST_ analysis of 56.4–57.2 Mb on chromosome 9, outlier loci in this region were colored in purple, the first-generation marker indicated in green and highlighted with a circle, and second-generation markers were highlighted with a circle. The tall plant *Dw1* gene at 57 Mb was noted with a black dashed line. **d**) Allele frequency of four second-generation markers on chromosome 9 in 30 Chinese and 390 SAP lines.

*Figure S5: Genotyping of segregants in the USDA-CSRL sorghum breeding program validated the functioning of second-generation markers in an independent breeding program*.

**a**) Second-generation marker S1_11126285 on chromosome 1 was used to genotype two intercross F_2_ populations. The F_2_ populations, generated by selfing the F1 progeny of BTx2752 × NSZ RIL and BTx642 × NSZ RIL, segregated in an expected ratio of 1:2:1 for XX:XY:YY (χ^2^ *p*-values 0.73 and 0.25). **b**) Second-generation marker S2_07404837 on chromosome 2 segregated the F_2_ populations BTx2752 × NSZ RIL and BTx642 × NSZ RIL in an expected ratio of 1:2:1 for XX:XY:YY (χ^2^ *p*-values 0.83 and 0.08).

*Figure S6: Seedling vigor1 comparisons of CT alleles in inbred NILs and hybrids*.

Performance of CT alleles in inbred near-isogenic lines (NILs) and hybrids carrying Chr2+, Chr4+, or −/− sib (no CT allele). Hybrids were generated by crossing five US elite R lines carrying the *male sterility3* (*ms3*) gene with inbred NILs. The US and Chinese parents were included as controls. Seedling vigor ratings showed an increase in inbreds NILs and hybrids with CT alleles compared to −/− sibs, however, no significant differences were observed between treatments.

## SUPPORTING TABLES

Supporting material available at Figshare: https://doi.org/10.6084/m9.figshare.21358191.v1

Table S1: KASP genotyping markers used for CT marker-assisted breeding with 100bp flanking sequences.

Table S2: Joint linkage mapping of NDVI values from AB17 field trial.

Table S3: Joint linkage mapping of NDVI values from MN17 field trial.

